# Conservation and divergence of DNA replication control in *Chlamydomonas reinhardtii*

**DOI:** 10.1101/2020.04.26.061929

**Authors:** Amy E. Ikui, Noriko Ueki, Kresti Pecani, Frederick Cross

## Abstract

We recently isolated temperature-sensitive cell cycle mutants in *Chlamydomonas reinhardtii* for which the causative mutations were located in genes annotated for potential involvement in DNA replication. *Chlamydomonas* has a very long G1 period during which cells grow up to ~10-fold without division, followed by rapid cycles of DNA replication and mitosis (‘multiple fission’). All of the candidate DNA replication mutants tested were defective in completion of the first round of DNA replication, and also failed to produce mitotic spindles. For a subset of the mutants, we rescued temperature-sensitive lethality with tagged transgenes and used the resulting strains to analyze abundance and localization control of the tagged protein. All of the DNA replication proteins tested were essentially undetectable until late G1, accumulated during the period of multiple fission and then were degraded as cells completed their terminal divisions. MCM4 and MCM6 were localized to the nucleus during the division cycle except for transient cytoplasmic localization during mitosis. CDC45 showed strict protein location to the nucleus and co-localized to spindles during mitosis. In contrast, CDC6 was detected in the nucleus only transiently during early divisions within the overall multiple fission cycle. Cdc6 protein levels were very low, but increased upon treatment with MG132, a proteasome inhibitor. We also tested if these DNA replication proteins are regulated by cyclin dependent kinase (CDK). There are two main CDKs in *Chlamydomonas*, CDKA1 and CDKB1. We found that CDC6 protein level was severely reduced in a *cdka1* mutant, but not in a *cdkb1* mutant. MG132 did not detectably increase CDC6 levels in the *cdka1* mutant, suggesting that CDKA1 upregulates CDC6 at the transcription level. Since MCM4, MCM6 and CDC6 were all essentially undetectable during the long G1 before DNA replication cycles began, we speculate that loading of origins with the MCM helicase may not occur until the end of the long G1, unlike in the budding yeast system. These results provide a microbial framework for approaching replication control in the plant kingdom.

## INTRODUCTION

The cell cycle is an ordered set of cellular processes in which DNA is replicated and segregated into two identical daughter cells. In eukaryotes, the cell cycle is controlled by the cyclin dependent kinase, CDK, coupled with cyclins [1]. *Chlamydomonas reinhardtii* is a single cell green algae which divides by ‘multiple fission’: a long G1 phase with massive cell growth, followed by typically 3-4 rounds of S/M phase that consists of DNA replication, chromosome segregation and cytokinesis [2]. The resulting daughter cells remain within the mother cell wall, finally hatching to produce 8-16 small newborn G1 cells that restart the cycle. There are two major CDKs in *Chlamydomonas reinhardtii*, *CDKA1* (a homologue of animal *CDK1/CDC2*) and *CDKB1* (a *CDK* specific to the plant kingdom) [3, 4]. While *CDK1* is the direct trigger of mitosis in yeast and animals, *CDKA1* in the plant kingdom specifically functions early in the cell cycle; mitosis is promoted by the plant-kingdom-specific *CDKB1*. One important role of *CDKA1* early in the cell cycle is transcriptional activation of a large battery of genes required for cell division that are turned on shortly before the division cycles begin [5, 6].

DNA replication has been studied extensively in the budding yeast *S. cerevisiae*. To initiate DNA replication in yeast, ORC (<u>O</u>rigin <u>r</u>ecognition <u>c</u>omplex) binds to origins followed by Cdc6 (<u>C</u>ell <u>d</u>ivision <u>c</u>ycle), Cdt1p and Mcm2-7 (<u>M</u>ini<u>c</u>hromosome <u>m</u>aintenance), forming the pre-replicative complex (pre-RC) [7–10]. CDK activity promotes export of the MCM complex from the nucleus, so that Mcm2-7 loading on chromatin is restricted to late M and early G1 phase when CDK activity is inhibited [11]. Firing of MCM-loaded (‘licensed’) replication origins, by contrast, requires CDK activity to phosphorylate multiple proteins to assemble a functional replication complex [12]. Once DNA replication is initiated, the pre-RC components such as Cdc6, Mcm2-7, and the ORC complex are phosphorylated by Cyclin/CDK; these multiple mechanisms prevent a second round of MCM reloading and consequent DNA replication from occurring before mitosis [13]. For example, Cdc6 is phosphorylated by Cyclin/CDK, targeting it for ubiquitin-mediated proteolysis through the SCF complex [14–17], Mcm2-7 is exported from the nucleus as noted above, and ORC function is inhibited by phosphorylation. These mechanisms collectively prevent origin relicensing within S phase.

The pre-RC proteins are mostly conserved in humans, but there are significant functional differences. While ScOrc1-6 binds to specific origin DNA sequences throughout the cell cycle, HsOrc1-6 is transiently associated with chromatin during G1 phase, and no origin consensus sequence has been defined [18]. HsMcm2 and HsMcm4 are phosphorylated by Cyclin B/Cdc2 during the G2-M phase [19]. Phosphorylation inhibits HsMcm2-7 reloading onto chromatin [20], but does not cause nuclear export as in yeast. Unlike in yeast, HsCdc6 is localized to the nucleus in G1 with a small fraction of HsCdc6 in the cytoplasm during S phase [21]. SCF-dependent HsCdc6 degradation inhibits DNA re-replication [22]. In contrast to yeast, where CDK-dependent phosphorylation promotes Cdc6 proteolysis, CDK-dependent HsCdc6 phosphorylation prevents Cdc6 proteolysis [23, 24].

After origin licensing, the pre-loading complex (pre-LC) is formed and activated by CDK and DDK kinases. Several critical proteins of the pre-LC in yeast are poorly conserved in vertebrates, such as Sld2. CDK-dependent ScSld2 phosphorylation is required for initiation of DNA replication [25, 26]. RecQL4 in vertebrates shows a unique N-terminal domain with limited homology to ScSld2, and a large C-terminal domain with 3’5’ DNA helicase activity. However, only the N-terminal region of RecQL4, lacking helicase activity and CDK phosphorylation sites, is required to trigger vertebrate DNA replication [39]. In yeast, Dpb11 is a scaffold protein used to recruit Sld2 and Sld3 through CDK dependent phosphorylation in order to assemble the Cdc45-Mcm-GINS (CMG complex). The human homologue of Dpb11, TopBP1, directly recruits Cdc45 onto replication origins [27]. Overall, the pre-LC is diverse both at the level of sequence and function between yeast and humans in contrast to the tight conservation of pre-RC across eukaryotes.

ORC, CDC6 and CDT1 have been identified in the land plant *Arabidopsis thaliana* [28]. AtORC1b is expressed only in proliferating cells. AtCDC6 is expressed during S-phase under control of E2F dependent transcription [29]. Ectopic expression of AtCDC6 or AtCDT1 induces endoreplication with altered ploidy levels [30]. AtCDC6 is degraded in a cell-free system in a proteasome-dependent manner [30]. AtMCM2-7 was expressed preferentially in tissues where DNA replication was occurring [31]. While AtMCM7 is transiently localized to the nucleus during G1 [32], AtMCM5 and AtMCM7 localize to the nucleus in G1, S and G2 phases and then translocate to the cytoplasm during mitosis [31]. MCM6 from maize is expressed at its highest level in the nucleus during G1 and then translocates to the cytoplasm during S phase [33]. Therefore, the function of MCM2-7 in plants may be regulated by localization. Plant *ORC* and *CDC6* have been shown genetically to be essential; however, the absence of conditional mutant alleles, and the lack of a simple cell cycle assay system in plants, has made it difficult to functionally address mechanisms of DNA replication control in the plant kingdom [34].

In this study, we analyzed DNA replication mutants identified in our previous UV-mutagenesis screening in *Chlamydomonas reinhardtii* [35, 36], concentrating on pre-RC components and their relationship to CDKA1 and CDKB1 activity.

## RESULTS

### Completion of the first cycle of DNA replication and nuclear division are inhibited in DNA replication mutants

In order to monitor cell cycle progression and to obtain a synchronous cell population in *Chlamydomonas reinhardtii*, we studied cells synchronized by nitrogen deprivation (causing arrest in G1) and refeeding. At intervals, cells were collected, fixed and stained with Sytox followed by flow cytometry analysis (FACS). By 12-14 hrs after release, wild type cells undergo multiple synchronous cycles of DNA replication and mitosis within the mother cell wall, yielding multinucleate cells producing signals of DNA content of 1, 2, 4, 8 and 16C (Figure 1A and 1B). Chloroplast DNA replication occurs early and uncoupled from the nuclear replication cycle and likely only accounts for a small part of the total FACS signal [37].

**Figure 1.**
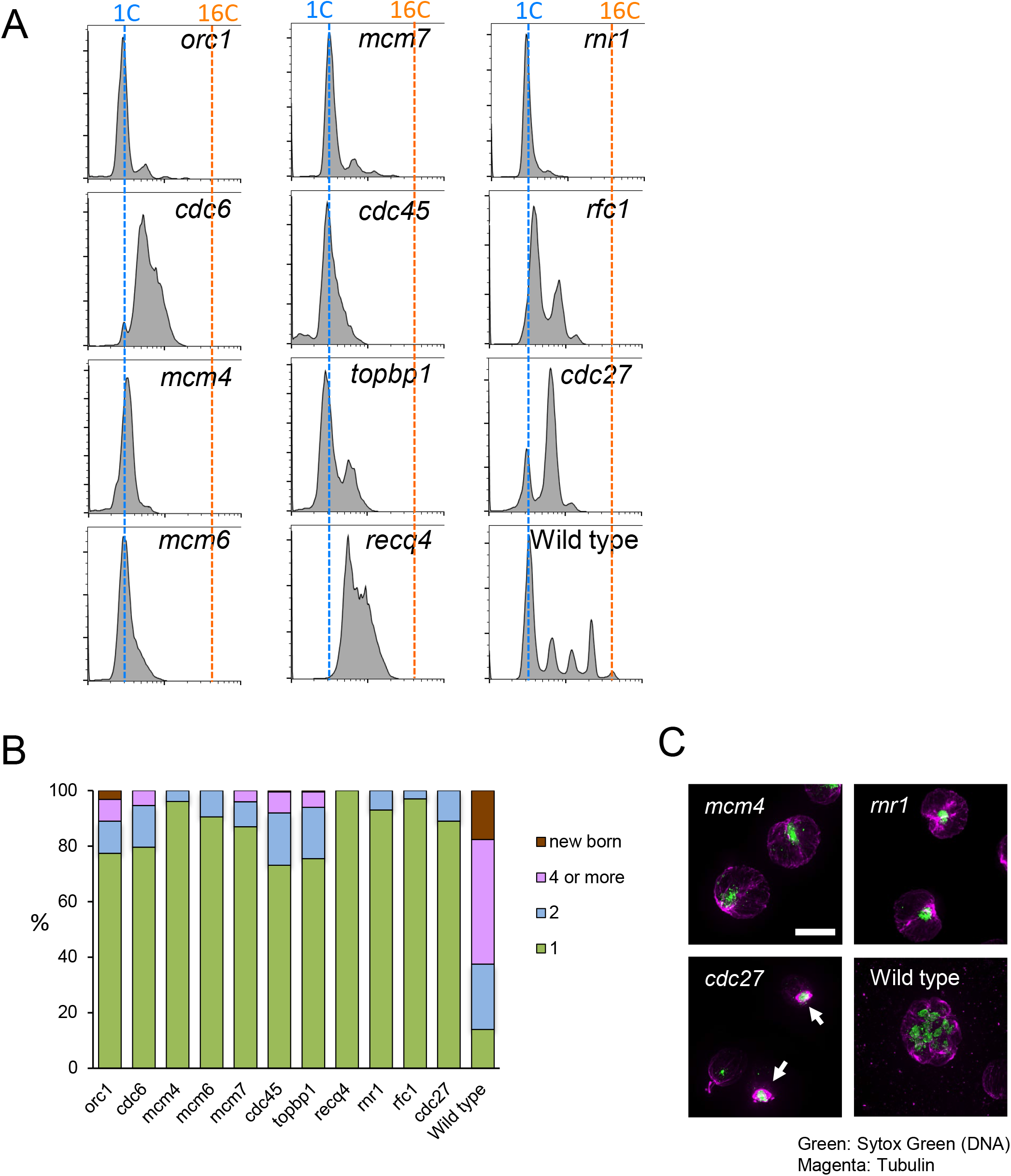
Analysis of DNA replication mutants (A) Wild type cells were arrested in G1 by nitrogen starvation and the cell cycle was released for 12hrs at the non-permissive temperatures 33°C. Cells were fixed and stained with Sytox for FACS. (B) Numbers of cells with new born, 4 or more nuclei, 2 nuclei or 1 nucleus was counted using the samples from A. 100 cells were counted and the percentage is shown. (C). Inhibition of spindle formations in DNA replication mutants. Indicated temperature sensitive mutants and Wild type cells were arrested in G1 and the cell cycle was released for 12 hours at 33 degrees. Cells were fixed and stained with anti-tubulin antibody (Magenta) and Sytox Green (Green). Images were taken by DeltaVision microscope. White arrows show mitotic spindles. Bars = 10 μm

The DNA replication mutant cells showed varying degrees of defect. Some showed no detectable increase in DNA content; others appeared to complete most of the first DNA replication cycle. However, all mutants were severely defective in nuclear division and entry into a second round of DNA replication (Figure 1A and 1B). Thus DNA replication is inhibited in the temperature sensitive (ts) DNA replication mutants as expected; in addition, incomplete DNA replication in the first cycle may block both nuclear division and reinitiation for a second round. Lack of nuclear division correlated with absence of detectable mitotic spindle formation by anti-tubulin immunofluorescence (Figure 1C). In yeast and animals, incomplete replication similarly triggers efficient blocks to mitotic progression [38].

### Amino acid sequence alignment of DNA replication proteins

Sequence alignments between the *Chlamydomonas* genes tested here generally showed the highest homology to land plant (*Arabidopsis*) genes, but also a strong alignment to animal and fungal sequences (Table 1). Interestingly, in many cases, alignment between *Chlamydomonas* and animal sequences was stronger than between *Chlamydomonas* and fungal sequences, most likely due to accelerated sequence evolution in fungal lineages. These alignments allow confident assignment of the *Chlamydomonas* mutants to the indicated orthologous animal sequences (Table 1).

**Table 1.**
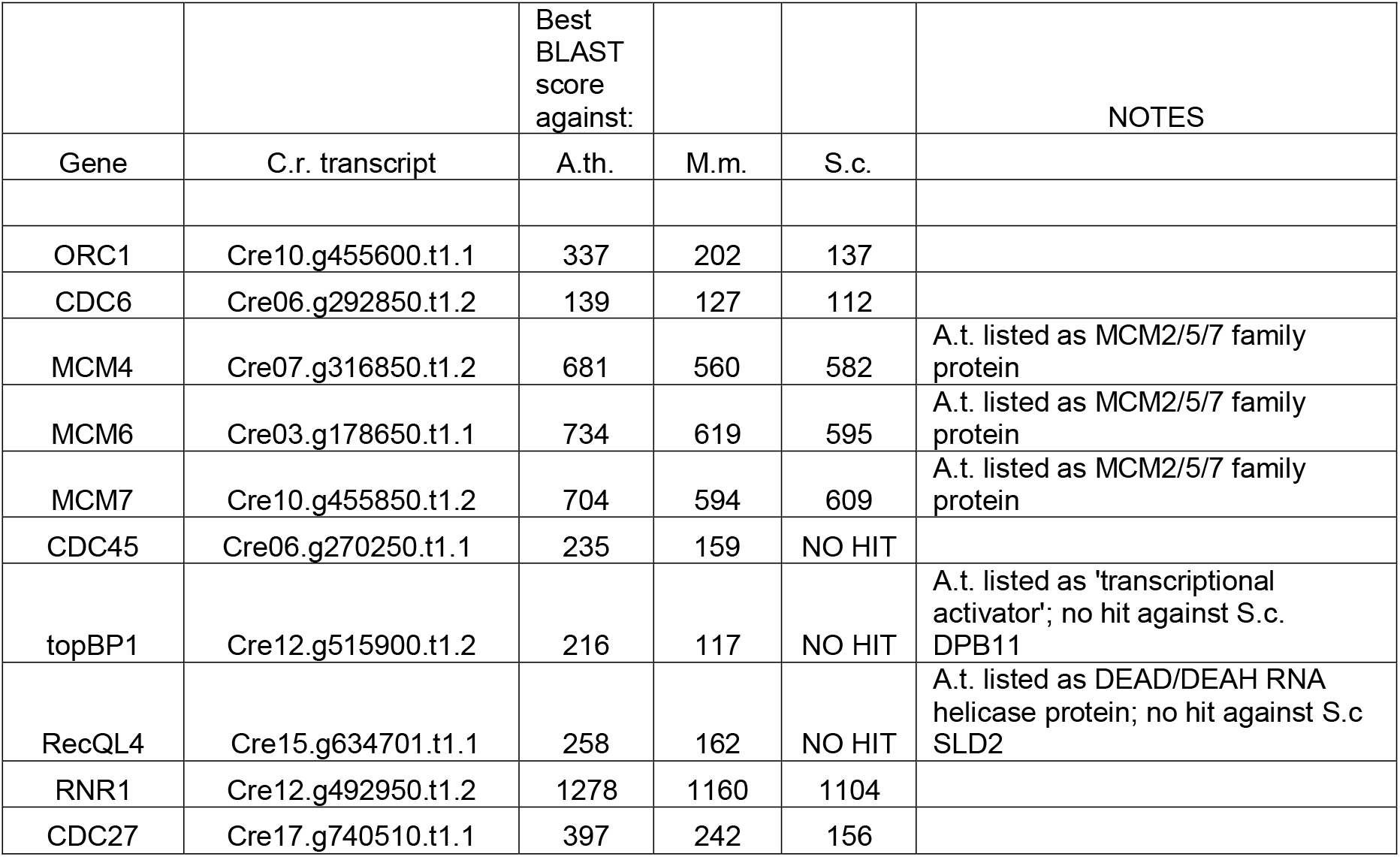

|  |  | Best BLAST score against: |  |  | NOTES |
| --- | --- | --- | --- | --- | --- |
| Gene | C.r. transcript | A.th. | M.m. | S.c. |  |
| ORC1 | Cre10.g455600.t1.1 | 337 | 202 | 137 |  |
| CDC6 | Cre06.g292850.t1.2 | 139 | 127 | 112 |  |
| MCM4 | Cre07.g316850.t1.2 | 681 | 560 | 582 | A.t. listed as MCM2/5/7 family protein |
| MCM6 | Cre03.g178650.t1.1 | 734 | 619 | 595 | A.t. listed as MCM2/5/7 family protein |
| MCM7 | Cre10.g455850.t1.2 | 704 | 594 | 609 | A.t. listed as MCM2/5/7 family protein |
| CDC45 | Cre06.g270250.t1.1 | 235 | 159 | NO HIT |  |
| topBP1 | Cre12.g515900.t1.2 | 216 | 117 | NO HIT | A.t. listed as 'transcriptional activator'; no hit against S.c. DPB11 |
| RecQL4 | Cre15.g634701.t1.1 | 258 | 162 | NO HIT | A.t. listed as DEAD/DEAH RNA helicase protein; no hit against S.c. SLD2 |
| RNR1 | Cre12.g492950.t1.2 | 1278 | 1160 | 1104 |  |
| CDC27 | Cre17.g740510.t1.1 | 397 | 242 | 156 |  |

The C-terminal region of Cre10.g455600 shared high-scoring alignment with MmOrc1 and also shared lesser homology with AtCDC6 and MmCdc6, confirming the similarity between Orc1 and Cdc6 [39]. ORC1 contains an ATPase associated with diverse cellular Activities (AAA+) domain, and the *div74-1* mutation was located in this domain (Figure 2A). Similar to MmOrc1, Cre10.g455600 also contains a PHD (<u>P</u>lant <u>h</u>omeo<u>d</u>omain) and BAH (<u>B</u>romo-<u>a</u>djacent <u>h</u>omology). Therefore, most likely, Cre10.g455600 is the ORC1 homologue in *Chlamydomonas*.

**Figure 2.**
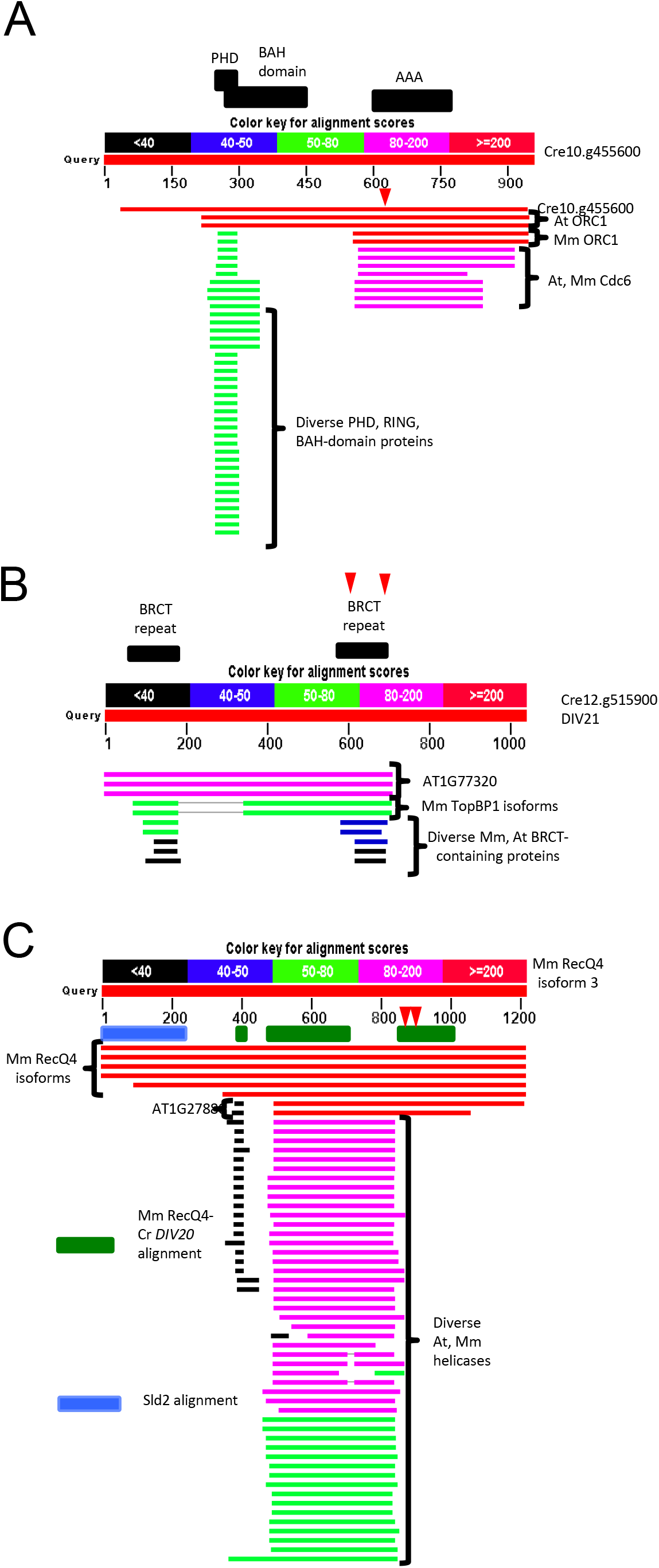
A graphical summary of sequence alignment by BLAST (A) *Chlamydomonas* Orc1 candidate, Cre10.g455600, was aligned with AtOrc1, MmOrc1, At Cdc6, MmCdc6 and proteins with PHD, RING and BAH domains. Red arrow indicates the location of *div74-1* mutation. (B) *Chlamydomonas* TopBP1 candidate, Cre12.g515900, was aligned with AT1G77320, MmTopBP1 and proteins with Mm, At BRCT domains. Black bar on the top shows BRCT repeat. Red arrows indicate the location of *div21-1* and of *div21-2* mutations. (C) Mm RecQ4 candidate was aligned with MmRecQ4 isofroms, AT1G2788, and At, Mm helicases. Blue bar shows Sld2 alignment. Green bar shows an alignment with *Cr DIV20* gene. Red arrows indicate the location of *div20-1* and of *div20-2* mutations.

As noted above, the functional equivalent of budding yeast Sld2 in animals is RecQL4. Homology between ScSld2 and animal RecQL4 is almost undetectable, at most, restricted to a small N-terminal region; however, this region, and not the large RecQL4 helicase domain, is required for DNA replication in *Xenopus* [40]. *Chlamydomonas DIV20* is a RecQL4 ortholog [35] that also has weak but detectable N-terminal ScSld2 homology (Table 1; Figure 2C). Surprisingly, though, mutations of two independent mutants that inactivate *DIV20* for DNA replication, map adjacent to the helicase domain in the C-terminal extension specific to the RecQ4 family. This region is dispensable for replication in *Xenopus* (Figure 2C) [40].

The RecQ family comprises a large number of proteins in *Arabidopsis* and other land plants. If our assignment of *DIV20* as a functional homolog of animal RecQL4 is correct, then we suggest that the *Arabidopsis* equivalent replication protein is AT1G27880 (Figure. 2C). Even though AT1G27880 is annotated (https://phytozome.jgi.doe.gov/) as a ‘DEAD/DEAH RNA helicase’, it is a protein with similar overall structure to *DIV20/*Cre15.g634701. Similarly, if *DIV21/Cre12.g515900* aligns functionally to animal TopBP1 for replication, then a likely *Arabidopsis* homolog might be AT1G77320, a BRCT-repeat-containing protein like TopBP1 and *Chlamydomonas DIV21* (Figure 2B). The functional significance of the BRCT repeats in *DIV21* is emphasized by the finding that two independent ts-lethal *div21* mutations lie in the region of one of its BRCT repeats. These examples point to the utility of functional identification of genes in the *Chlamydomonas* genome (mainly single-copy) to help navigate the more complex, geneduplicate-rich land plant genomes.

### Synthetic lethality between pre-RC mutants

If *Chlamydomonas* pre-RC components function in complexes, as do their yeast and human orthologs, then strong genetic interactions might be expected between mutations in different genes in this set. We inter-crossed pre-RC mutants *orc1, cdc6, mcm4, mcm6-981A* and *mcm6-GHI* [36] and analyzed viability of double mutant progeny at the permissive temperature. *mcm6-981A* was synthetically lethal with *orc1, cdc6* and *mcm4* (Figure 3A). *mcm6-GHI*, showed mild synthetic lethality with *orc1*, though it was not lethal at the permissive temperature when combined with *cdc6* (Figures 3B). The *orc1 cdc6* double mutant was also mildly sick (Figure 3C). The genetic interaction was summarized in Figure 3D. Frequent synthetic lethality in this mutant set is consistent with the proteins that form functional complexes, as expected from prior work in yeast (see Introduction).

**Figure 3.**
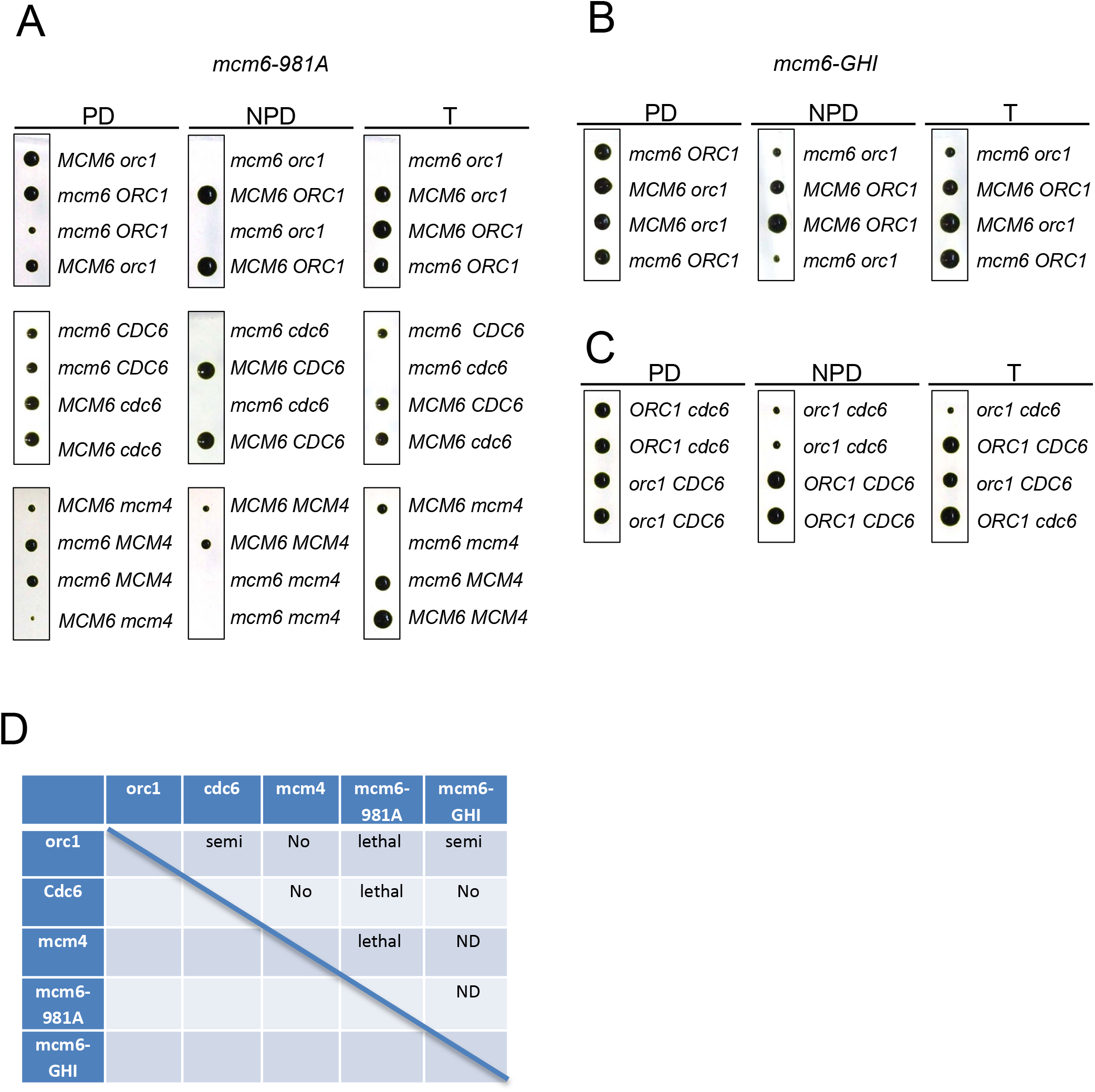
Synthetic lethality between DNA replication mutants (A) *mcm6-981A* temperature sensitive mutant was crossed with *orc1, cdc6* or *mcm4* mutant. Tetrads are shown with tetra type. An example of parental ditype (PD), non-parental ditype (NPD) or tetra type (T) is shown. (B-C) Tetrad analysis of *mcm6-GHI orc1* and *orc1 cdc6* are shown. (D) A summary table of synthetic lethality tested in this study. Lethal: complete lethal, Semi: mild lethality, No: No genetic interaction, ND: No data.

### Transgenes expressing fluorescent fusion proteins rescued temperature sensitivity of the replication mutants

We created universal tagging plasmids containing mCherry, Venus or GFP to be used in *Chlamydomonas*. The plasmid backbone was generated from *CDKB1* plasmid which contains mCherry-tagged *CDKB1* gene [41]. The final plasmids contain *aphVIII CDS* for paramycin resistance that is useful for drug selection in *Chlamydomonas*, an Ampicillin resistant gene for bacterial selection, and mCherry, GFP or Venus followed by the *CDKB1* 3’UTR, as well as multiple cloning sites for insertions of desired promoters and coding sequences (Figure 4A).

**Figure 4.**
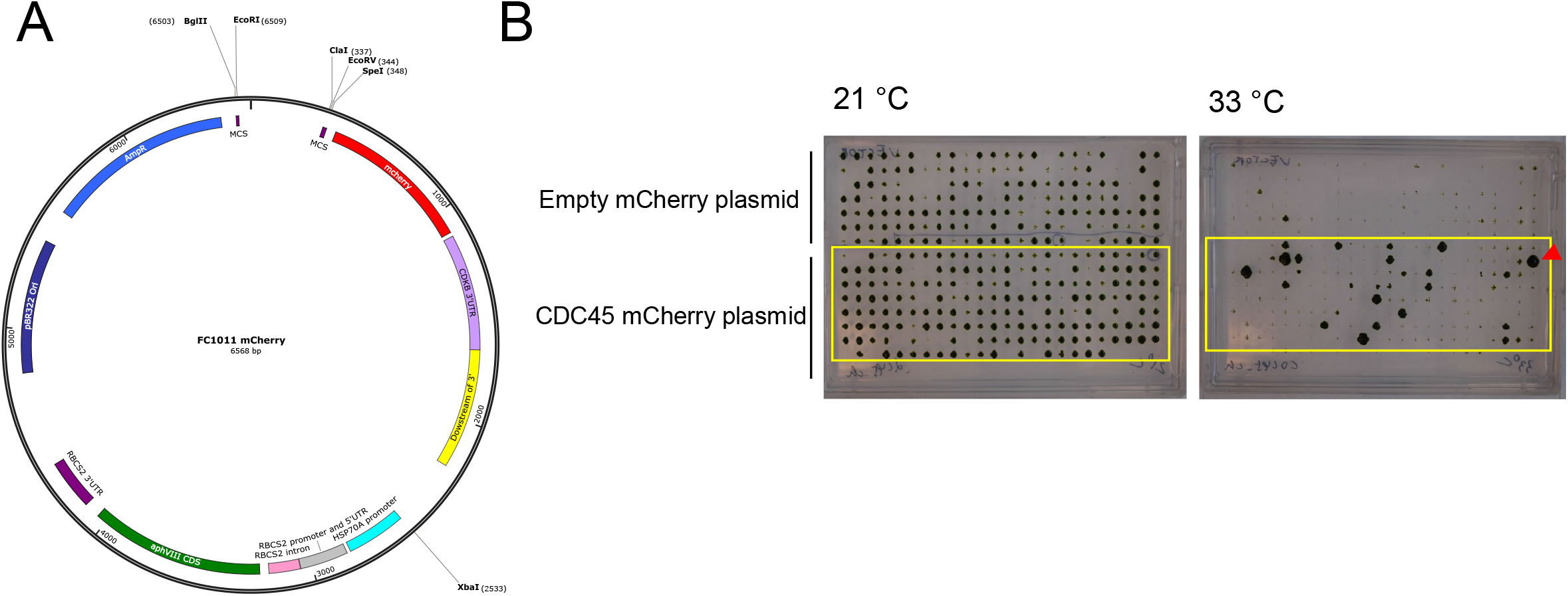
Chlamydomonas plasmid construction. (A) An universal mCherry plasmid map is shown. Tagging can be mCherry, Venus or GFP. (B) CDC45-mCherry plasmid was constructed and transformed into *cdc45* temperature sensitive mutant. Transformants were picked from plates at 21 °C and aligned as 364 well format. The plates were incubated at 21 or 33 °C for 10 days. The colony with red arrow was further analyzed.

We amplified *CDC45* PCR product from genomic DNA and inserted it into to the mCherry plasmid through Gibson assembly. The resulting *CDC45-mCherry* plasmid was linearized with XbaI and electroporated into the *cdc45* ts mutants. Tranformants were selected for antibiotic resistance at 21°C (permissive temperature) and then screened at the non-permissive temperature, 33°C (Figure 4B). A small proportion of transformants grew at 33°C; such low co-rescue is a general feature of *Chlamydomonas* transformation, probably resulting from frequent breakage or silencing of transforming DNA. Colonies that grew at 33°C were picked and their genomic DNA was tested by PCR for the presence of mCherry, the region covering the promoter and the ampicillin resistance gene. No temperature-resistant colonies were obtained with the empty vector (Figure 4B).

Similarly, we constructed tagging plasmids to rescue other ts-lethal mutants. Efficiency of rescue in the selected transformants was confirmed by serial dilution (Figure 5A). Selected transformants were further subjected to western blotting to analyze the expression and size of the fusion protein (Figure 5B). Clones with efficient rescue and protein expression were backcrossed with wild type at least once before use in further experiments (Figure 5, red numbers). Transgenes in *Chlamydomonas* integrate at random locations, which could affect expression efficiency. In addition, the transgene plasmid is often broken or rearranged in *Chlamydomonas* before it is integrated. The Western blot analysis and genetic rescue ensures an approximately full-length fusion protein, but in principle the promoter region could be disrupted causing less efficient transcription. In order to maximize the likelihood that the selected transformant would contain a complete transgene, we designed the primers to detect inclusion of the promoter, ampicillin resistant gene and inclusion of the tagged gene.

**Figure 5.**
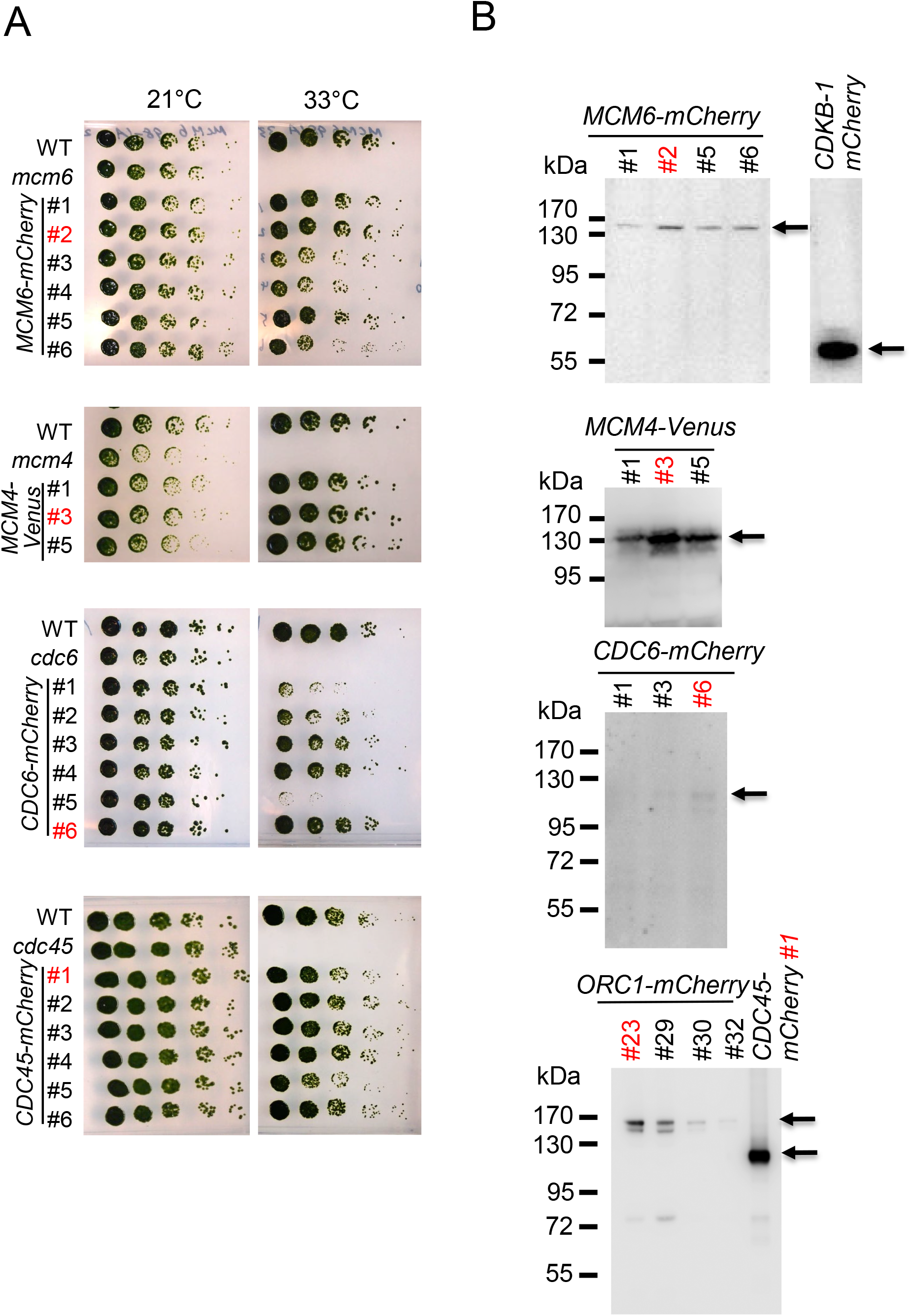
Transgene of *MCM6-mCherry, MCM4-Venus, CDC6-mCherry, CDC45-mCherry* rescued temperature sensitivity of the corresponding mutant. (A) Indicated strains were grown and serially diluted 5fold on TAP plates. The plates were incubated at 21 or 33 °C for 5-7 days. The strain number in red was used in the following experiments. (B) Protein expression of MCM6-mCherry, MCM4-Venus, CDC6-mCherry, ORC1-Venus and CDC45-mCherry were detected by western blotting using anti-mCherry or anti-GFP antibody. CDKB 1-mCherry is shown as a positive control.

### Expression of DNA replication proteins during the division cycle

All transgenes were constructed with their native promoters, so that we could analyze accumulation of MCM6-mCherry, CDC45-mCherry, ORC1-Venus and CDC6-mCherry fusion proteins at endogenous levels throughout the cell cycle. We included a strain expressing CDKB1-mCherry as a positive control since it is known to accumulate specifically in division-phase cells [41]. Cells were arrested at G1 phase as described above, then released into complete media at 33°C. Since the transgene was expressed in the context of the endogenous ts-lethal mutation, cell cycle progression was dependent on the fluorescent fusion protein in each case. CDKB1-mCherry was first expressed at 10 hrs, maximized around 14 hrs and declined by 24 hrs as previously shown [41]. These times correlate with the period of multiple fission cycles. Similarly, the maximum levels of tagged MCM6, ORC1, CDC6 and CDC45 were observed at 14 hrs after release (Figure 6A and 6B). ORC1 and CDC6 mCherry fusion protein levels were undetectable by direct Western analysis. Therefore these tagged proteins were pulled down using Dynabeads conjugated with mCherry nanobody to concentrate the protein before Western analysis. FACS analysis of *CDC6-mCherry* showed that DNA replication initiated at 12 hrs as expected in this procedure (Figure 6C). We conclude that pre-RC proteins are essentially absent during the long G1 phase in *Chlamydomonas*, and like CDKB1 are expressed only during the rapid divisions of the multiple fission cycle. This is an interesting contrast to the yeast system, in which pre-RC components are loaded onto origins starting in late mitosis and are bound throughout G1 phase.

**Figure 6.**
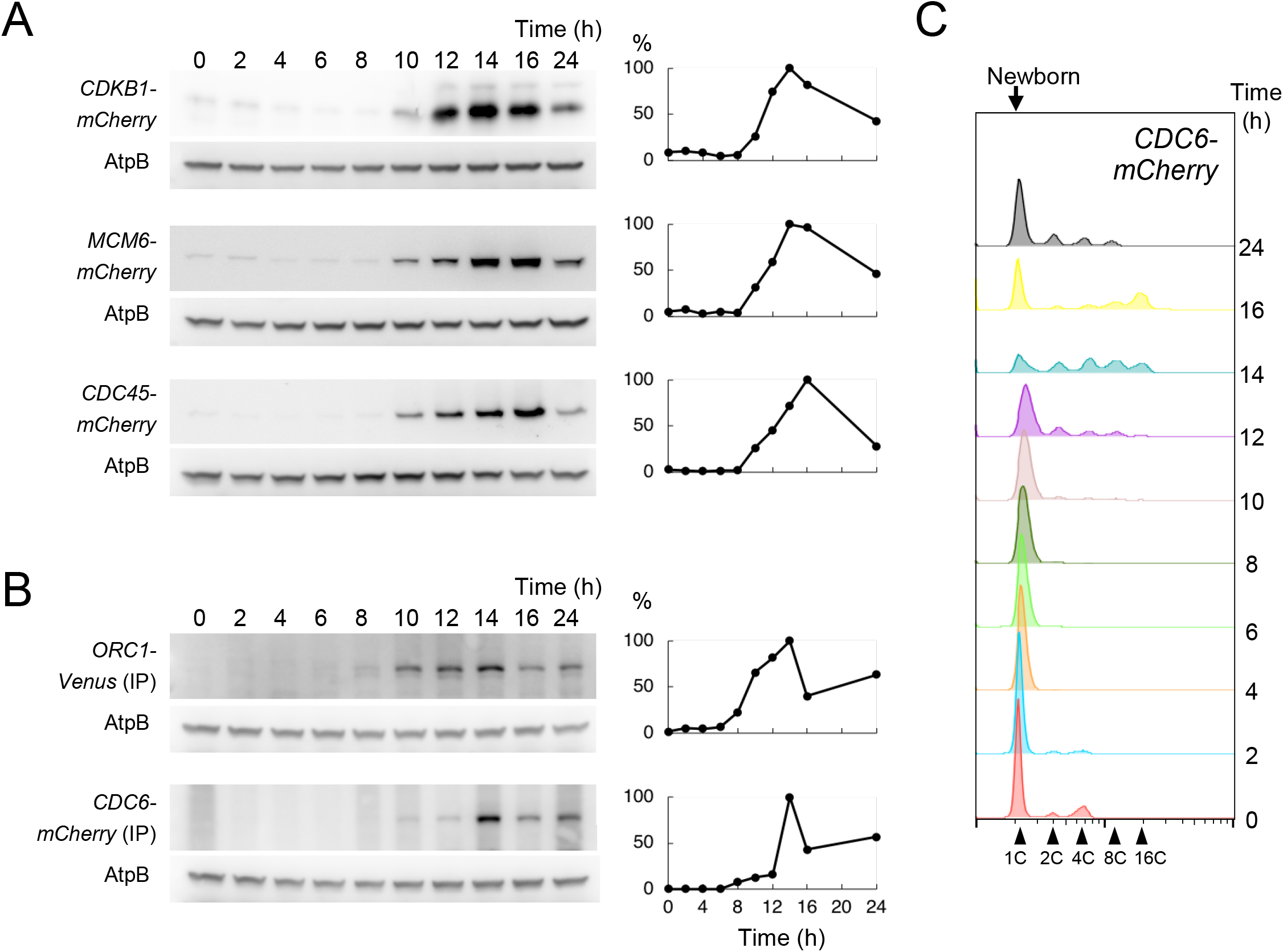
DNA replication proteins are expressed during cell division. (A) *CDKB-mCherry, MCM6-mCherry* or *CDC45-mCherry* cells were synchronized in G1 by nitrogen starvation. The cell cycle was released and cells were collected at indicated time points. Protein was extracted and subjected to western blotting analysis. The signal was detected by anti-mCherry antibody. AtpB was used as a loading control. The relative protein expression quantified by ImageJ is shown on the right. The highest expression during the cell cycle was set as 100%. (B) *ORC1-Venus* or *CDC6-mCherry* cells was prepared as described in A. Protein was pulled down by GFP or mCherry nanobody coupled with DynaBeads. The signal was detected by anti-GFP or anti-mCherry antibody. AtpB was used as a loading control. The relative protein expression is shown in the right. The highest expression during the cell cycle was set as 100%. (C) The *CDC6-mCherry* samples used in B were fixed, stained and subjected to FACS analysis to monitor cell cycle progression.

### MCM proteins are localized to the nucleus during the division cycle with a transient release during mitosis

Mcm6-mCherry protein localization was analyzed by indirect-immunofluorescence using anti-mCherry antibody. Mcm6 was localized to the nucleus in wild type cells, though sporadic cells with dispersed Mcm6 protein in the cytoplasm were detected (Figure 7A, white triangles). We used a *cdc27* cell cycle mutant in order to examine Mcm6 localization in metaphase-blocked cells. CDC27 is a part of the <u>A</u>naphase <u>P</u>romoting <u>C</u>omplex (APC) [42], required for the metaphase-anaphase transition [35, 41]. MCM6 was diffused from the nucleus in metaphase-arrested *cdc27* cells, and co-staining of microtubules showed that these cells contained mitotic spindles (Figure 7B). After 16 hrs the cell cycle was completely arrested in G2/M phase after completion of a single round of DNA replication in the *cdc27* mutant (Figure 7B). Most likely, the sporadic wild type cells with diffuse MCM6-mCherry localization are cells transiting through metaphase.

**Figure 7.**
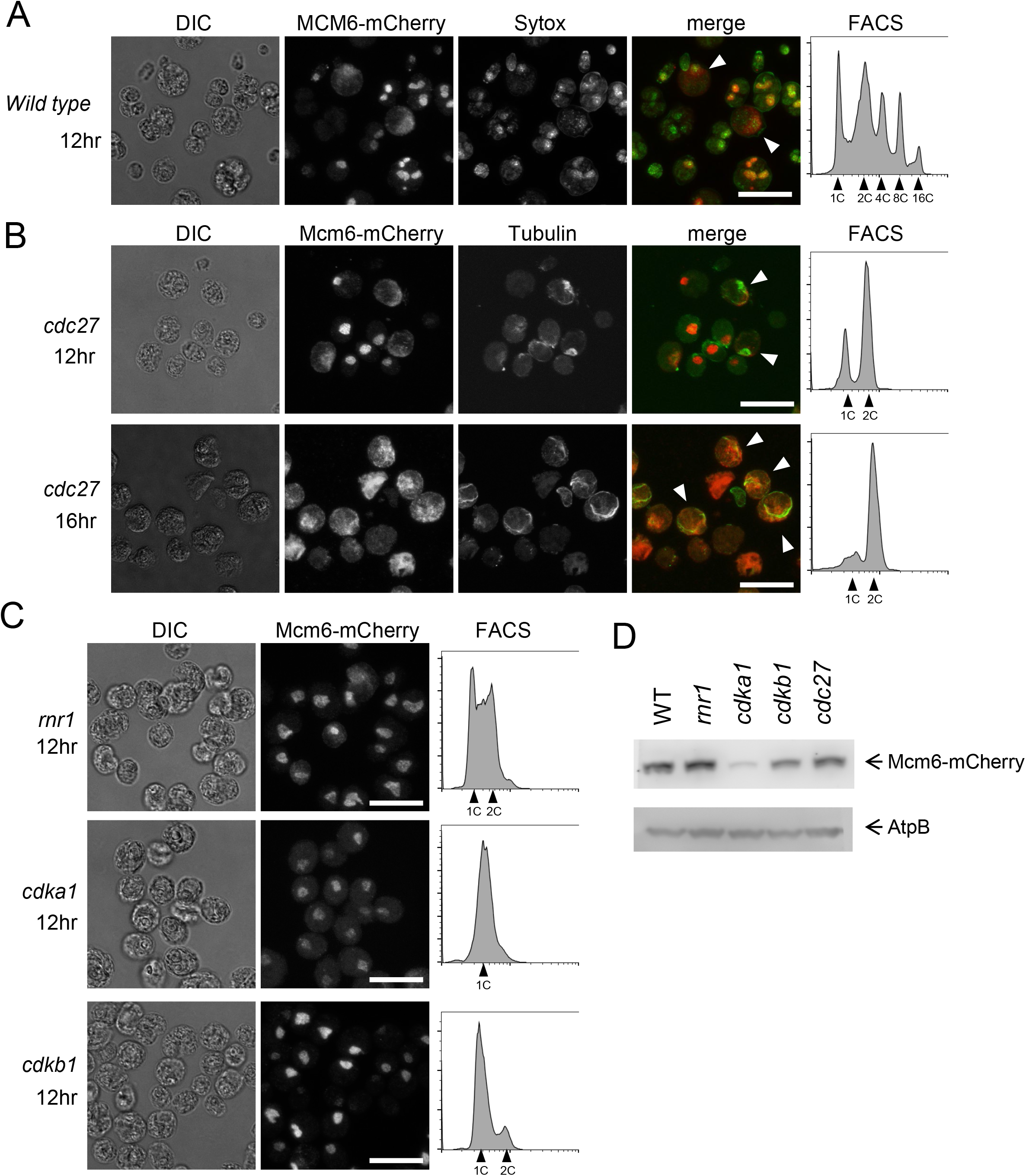
MCM6 is localized to the nucleus in dividing cells. (A) *MCM6-mCherry* cells (Wild Type) were arrested in G1 and released at 33 °C for 12hrs. Cells were fixed and stained with anti-mCherry antibody (Red) and Sytox green (Green). White triangles indicate cells with diffused Mcm6. Images were acquired by Confocal microscope. Bars = 20 μm. The DNA content was analyzed by FACS (right panel) (B) *MCM6-mCherry cdc27* cells were collected after 12hrs or 16hrs from release and prepared as described in A. White triangles show cells with mitotic spindles. Bars = 20 μm. The DNA content was analyzed by FACS (right panels). (C) *MCM6-mCherry rnr1, MCM6-mCherry cdka1* or *MCM6-mCherry cdkb1* cells were collected after 12hrs from the release. Cells were prepared as described in A. Bars = 20 μm. The DNA content was analyzed by FACS (right panels). (D) Indicated strains were arrested in G1 and collected 12hrs after the release. Proteins were extracted and MCM6 was detected by anti-mCherry antibody. AtpB was used as a loading control.

In budding yeast, translocation of Mcm2-7 to the cytoplasm in S-phase after origin firing may prevent origin re-loading within a given S-phase [43]. Localization of Mcm2-7 is controlled by Cdk1-dependent phosphorylation [44, 45]. It is possible that transient mitotic loss of MCM6 from the nucleus in *Chlamydomonas* is regulated similarly. We tested if CDKA1 or CDKB1 controls MCM6 localization and found that MCM6 stayed in the nucleus in both *cdka1* and *cdkb1* mutants (Figure 7C), while leaving the nucleus in the *cdc27* background in which CDKA1 and CDKB1 activities are both high [48]. Thus it is possible that CDK activity regulates Mcm6 localization in *Chlamydomonas* as in yeast; we lack direct evidence for this idea, however. Western blotting analysis showed that the MCM6-mCherry protein was readily detectable in *rnr1, cdkb1* and *cdc27* mutants similar to the level in wild type cells and was slightly suppressed in the *cdka1* mutant (Figure 7D).

To extend these findings to a different member of the MCM complex, we monitored MCM4-Venus localization in single cells, using a variant of procedures developed for budding yeast time-lapse microscopy [46, 47] (see Methods). Cells expressing MCM4-Venus were monitored at 3-min resolution, starting in G1 and extending to completion of a multiple fission cycle. MCM4-Venus was absent in the initially plated G1 cells, but began to accumulate approximately 2 hrs before the first division (Figure 8A) (Movie1). MCM4-Venus was localized to a small region in the cell anterior that we assume is the cell nucleus based on size, shape, position and similarity to the MCM6-mCherry nuclear localization in immunofluorescence experiments in Figure 7. The MCM4-Venus signal was greatly reduced from the nucleus in the frame right before the first division, but reappeared almost immediately after division (Figure 8A and 8C). Quantification of the total MCM4-Venus signal across the cell indicated that the reduction or loss of nuclear intensity occurred without significant change in total cellular levels. Therefore, we attribute the loss of nuclear signal to transient diffusion into the cytoplasm rather than to MCM4 proteolysis and rapid resynthesis. MCM4-Venus transient loss was restricted to a single 3-min frame in almost all cells observed. We observed the same one-frame loss of MCM4-Venus nuclear intensity right before the second and third divisions, again with no notable change in total cellular MCM4-Venus levels (Figure 8A and 8C). As with the first division, MCM4-Venus nuclear signal returned immediately on cytokinesis at completion of the later divisions. In the *cdc27* background, MCM4-Venus was localized in the nucleus in G1, but delocalized as cells entered the *cdc27* arrest, unlike in wild type, where MCM4-Venus delocalization lasted for only ~3min coincident with cell division. MCM4-Venus remained diffuse over many hours (Movie2) in the *cdc27* background. Overall, these results indicate a metaphase dispersal of MCM4, just as was observed with MCM6 above, and suggest that the entire MCM complex may be transiently delocalized from the nucleus to the cytoplasm at metaphase.

**Figure 8.**
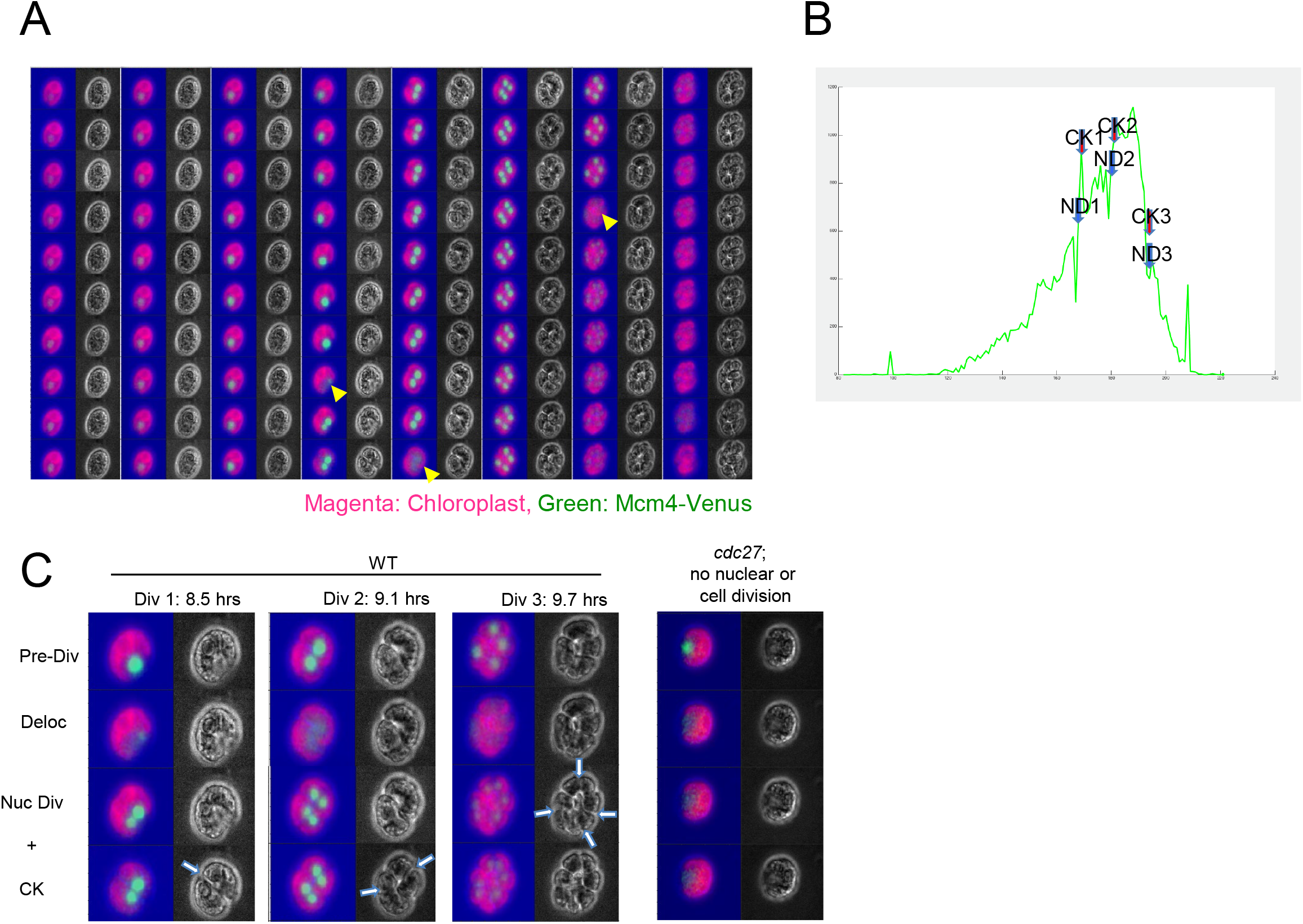
MCM4 is briefly translocated to the cytoplasm in every single S/M phase. (A) *MCM4-Venus* cells were blocked in G1 by nitrogen starvation and placed on TAP for live imaging by time-lapse microscopy. MCM4-Venus (Green) and chloroplast auto-fluorescence (Magenta). Yellow arrows indicate the time when Mcm4-Venus disappeared from nucleus. (B) The relative intensity of MCM4-Venus signal in (A) was quantified. ND: nuclear division, CK: cytokinesis. (C) The representative images of four stages (Pre-Div: pre division, Deloc: delocalization, Nuc Div: nuclear division, CK: cytokinesis) are shown. Left columns: 1st, 2nd and 3rd division in wild type, right column: 1st division in *cdc27*. Blue arrows indicate the division site.

To determine if this transient protein delocalization in metaphase is specific to the MCM proteins, we used a strain expressing a bleomycin-resistance gene fused to GFP (Ble-GFP). Unlike free GFP, Ble-GFP is nuclear-localized [48], presumably due to nuclear targeting of Ble. Much as with MCM4-Venus, Ble-GFP nuclear signal was sharply reduced for about 3min right at the time of cell division, with no decrease in total cellular Ble-GFP signal (Movie 3). Ble-GFP in the *cdc27* background diffused from the nucleus as cells entered a *cdc27* arrest and failed to re-concentrate for the duration of the movie (Movie4). Thus, Ble-GFP reconstitutes the cell-cycle-regulated nuclear-cytoplasmic traffic observed with MCM4 and MCM6.

We were unable to analyze ORC1-Venus and CDC6-Venus by time-lapse microscopy, presumably due to their low abundance as noted above from Western analysis in Figure 6B. However, cytoplasmic diffusion of CDC6-mCherry was observed by indirect immunofluorescence in the *cdc27* mutant, instead of nuclear concentration observed in wild type (Figure 9B; see below).

**Figure 9.**
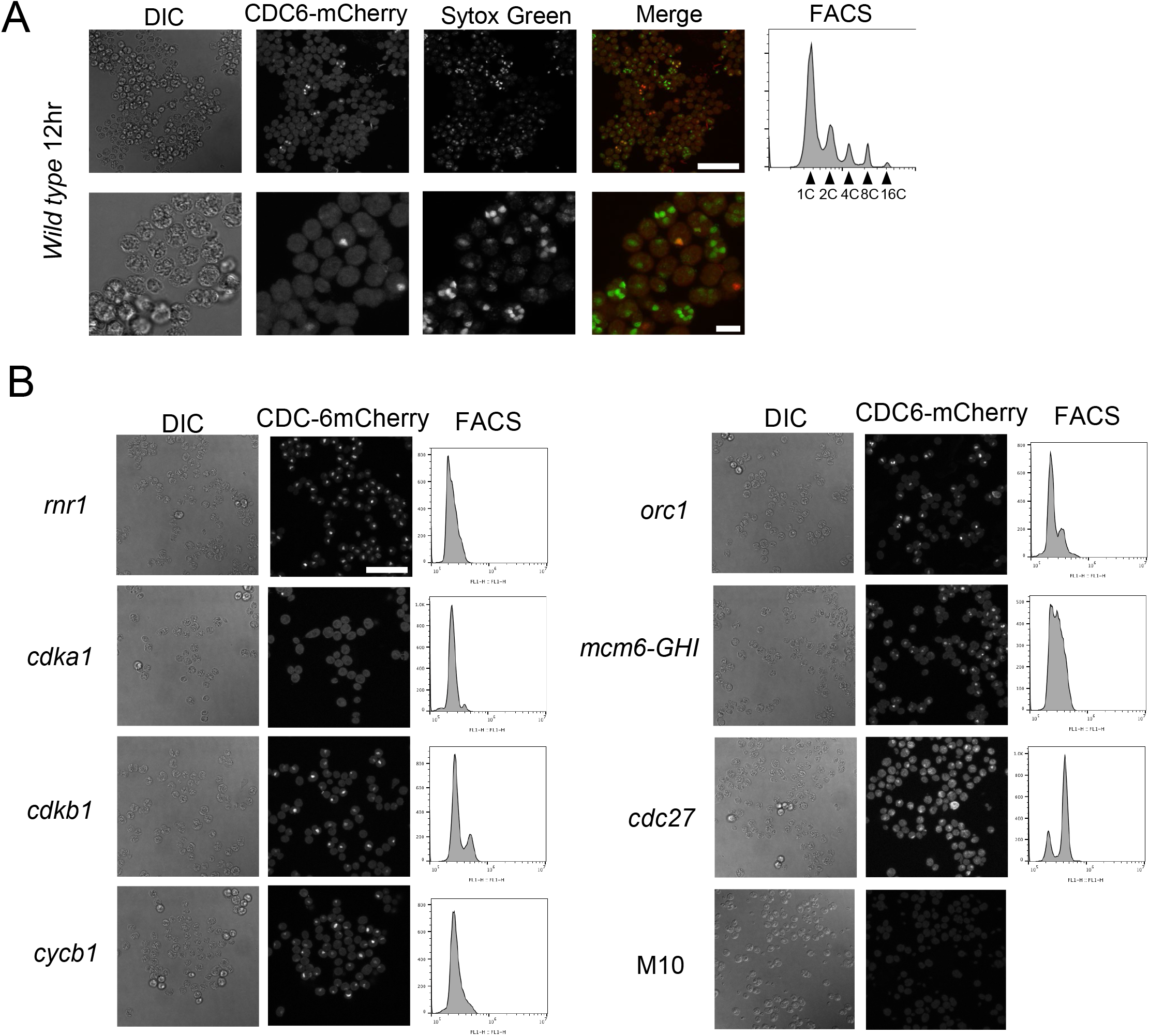
CDC6 protein is transiently localized to the nucleus in dividing cells. (A) *CDC6-mCherry* (Wild Type) cells were synchronized in G1 and collected 12 hours after the release. Cell were fixed and stained with anti-mCherry antibody (Red) and Sytox green (Green). Images were acquired by Confocal microscope. Scale bar = 50 μm (upper row) and 10 μm (lower row). DNA content was analyzed by FACS (right panel). (B) *CDC6-mCherry* in *rnr1, cdka1, cdkb1, cycb1, orc1, mcm6-GHI* or *cdc27* mutant was synchronized in G1 and collected 14 hrs after the release. Cell were fixed and stained with anti-mCherry antibody. Images were acquired by Confocal microscope. DNA content was analyzed by FACS (right panel). M10 strain without *CDC6-mCherry* was used as a negative control to show background signal. Scale bar = 50 μm.

### CDC6 protein levels are dependent on CDKA1 activity

By immunofluorescence, CDC6-mCherry was detectable in cell nuclei, although almost never in cells with more than two nuclei. This did not reflect completion of cell divisions, since these cells, in general, underwent 3-4 division cycles (yielding 8-16 nuclei) (Figure 9A). The model from budding yeast (see Introduction) suggests that CDC6 should be required at the end of each mitosis to license replication for the succeeding rounds. We don’t know if this is a real difference between the systems or if late rounds of replication in *Chlamydomonas* can be licensed with very low (undetectable) levels of Cdc6.

In budding yeast, Cdc6 protein is regulated by transcription, localization and protein degradation (see Introduction). Cdc6 is phosphorylated by Cdk1 and targeted for ubiquitination followed by proteasome-mediated protein degradation [14–16, 49–51]. Therefore, we tested if *Chlamydomonas* CDC6-mCherry protein levels are affected by inactivation of *CDKA1* or *CDKB1*. CDC6-mCherry was transiently localized to the nucleus during the division cycle in wild type cells, and also nuclear-localized in *rnr1, cdkb1, cycb1, orc1* and *mcm6-GHI* mutants (Figures 9A and 9B). By contrast, in the *cdka1* mutant, CDC6-mCherry was completely undetectable (Figure 9B). Thus CDKA1 is required for efficient CDC6 accumulation. Unlike in yeast, where CDK activity results in destabilization and lower CDC6 levels, in *Chlamydomonas* the reverse is observed.

To confirm and extend these findings, we examined CDC6-mCherry protein level by western blot in various mutant backgrounds. Results were consistent with the immunofluorescence results: CDC6-mCherry was observed in *rnr1, cdkb1, cycb, orc1, mcm6-981A, mcm6-GHI* and *cdc27* after 12 hrs from release (Figure 10A), but was undetectable in the *cdka1* mutant at 12 and 14 hrs after release (Figure 10A). This finding was reproduced with three independent *CDC6-mCherry cdka1* clones, where in all cases the *CDC6-mCherry* transgene efficiently rescued ts–lethality of the *cdc6-1* mutation (data not shown).

**Figure 10.**
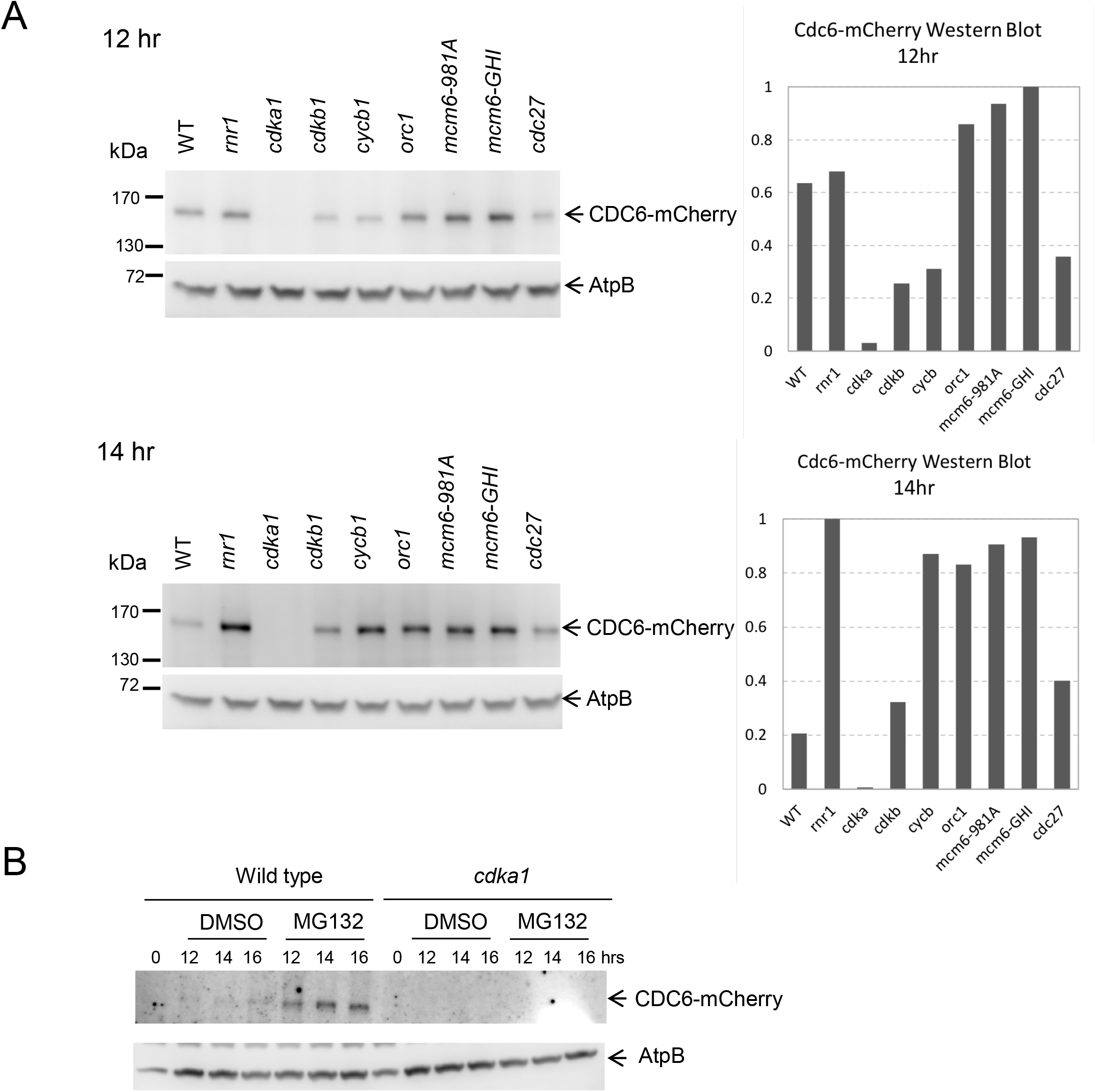
CDC6 protein expression in cell cycle mutants (A) CDC6 protein expression was suppressed in *cdka1* mutant. Indicated strains were arrested in G1 and collected 12hrs or 14hrs after release. Proteins were extracted and CDC6-mCherry was pulled down using mCherry nanobody coupled with DynaBeads. The signal was detected by anti-mCherry antibody. AtpB was used as a loading control. The graph shows quantification of relative expression in each strain. (B) *CDC6-mCherry* (WT) or *CDC6-mCherry cdka1 (cdka1*) cells were arrested in G1 and plated on TAP plates to release the cell cycle arrest. After 10hrs from release, cells were plated on TAP with DMSO or MG132. Cells were collected 12, 14 or 16hrs after release corresponding to 2, 4 or 6hrs treatment. Proteins were extracted and CDC6-mCherry was pulled down using mCherry nanobody coupled with DynaBeads. The signal was detected by anti-mCherry antibody. AtpB was used as a loading control. The graph shows quantification of relative expression in each strain.

*CDC6* is still essential in a *cdka1* background, since *cdka1;cdc6-1* double mutants are temperature-sensitive but *cdka1;CDC6* single mutants are not (data not shown). This finding implies that there must be an undetectable but still functional level of CDC6 expressed in the *cdka1* mutant. The prolonged delay in replication in *cdka1* mutants [5] may be due at least in part to very low levels of this essential replication protein.

### *Chlamydomonas* CDC6 is degraded by the proteasome

In yeast, CDK-phosphorylated Cdc6 is degraded by ubiquitin-mediated proteolysis [14–16, 49, 51]. We examined *Chlamydomonas* CDC6 accumulation in the presence of a proteasome inhibitor, MG132. Cells were released from G1 and collected after 12, 14 or 16 hrs. In wild type cells, CDC6 levels were increased by MG132 treatment (Figure 10B), suggesting that CDC6 is degraded by the proteasome. However, MG132 treatment did not result in detectable CDC6 accumulation in the *cdka1* background (Figure 10B). Therefore, CDC6 suppression in the *cdka1* mutant is unlikely to be due solely to excessive proteasomal degradation. In animal cells, MG132 blocks cells in metaphase. However, in our experiments, the *Chlamydomonas* cell cycle profile was not significantly changed by treatment (data not shown). We do not know if failure of MG132 to arrest the *Chlamydomonas* cell cycle is due to inefficient proteasome inhibition at the levels we used or for some other reason. However, the cell cycle progression in MG132-treated cells does allow us to conclude that the CDC6 stabilization we observe is not due to an indirect effect of cell cycle arrest.

### CDC45 is nuclear and may interact with mitotic spindles

CDC45 is a key protein required for conversion of a replication origin loaded with Mcm2-7 to a firing replication fork [12]. Cdc45-mCherry localization was analyzed by indirect immunofluorescence. We found that CDC45-mCherry was localized to the nucleus (Figure 11A). However, there was no transient CDC45 diffusion observed in any cells. In the *cdc27* mutant, CDC45-mCherry stayed in the nucleus and was co-localized with tubulin (mitotic spindle) staining (Figure 11B). We conclude that CDC45 was not diffused to the cytoplasm at metaphase, unlike MCM4, MCM6, CDC6, and ble-GFP (see above). This may be due to tethering of CDC45 to the mitotic spindle. The function of CDC45 interaction with the mitotic spindle is unknown.

**Figure 11.**
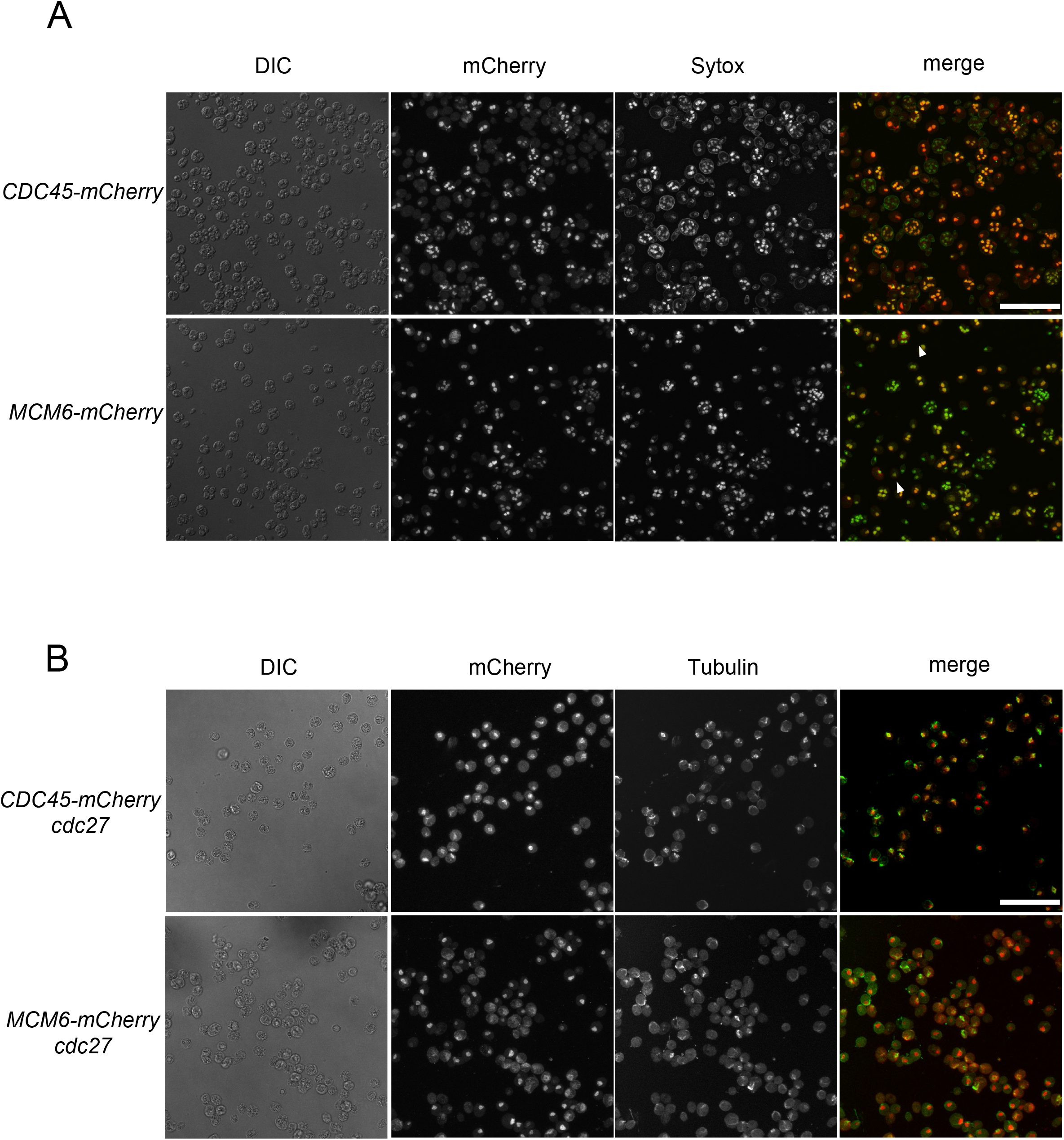
Cdc45 protein is localized to spindles. (A) *CDC45-mCherry or MCM6-mCherry* cells were synchronized in G1 and collected 12hrs after release. Cell were fixed and stained with anti-mCherry antibody (Red) and Sytox green (Green). Images were acquired by Confocal microscope. Scale bar = 50 μm. (B) *CDC45-mCherry cdc27* or *MCM6-mCherry cdc27* mutant was synchronized in G1 and collected 12 hrs after release. Cell were fixed and stained with anti-mCherry or anti-tubulin antibody. Images were acquired by Confocal microscope. Scale bar = 50 μm.

## DISCUSSION

### A conserved set of replication control proteins in the plant kingdom

Our previous hunt for high-throughput temperature-sensitive mutants generated hundreds of cell cycle mutants, for which the causative mutations were determined by multiplexed bulked-segregant sequence analysis [35, 36]. A number of these mutations fell in proteins aligning with replication control proteins defined in other eukaryotes. We show here that these mutants were indeed defective in DNA replication, indicating that their core functions as well as their sequences are conserved (Figure 1).

Inhibition of replication by any of these mutants also blocked mitotic spindle formation (Figure 1C). This could reflect a checkpoint preventing mitotic entry until completion of replication as observed in many other eukaryotes [38]. A checkpoint of this kind may thus be conserved in *Chlamydomonas*, and probably in the plant kingdom overall [52]. In contrast, another well-known checkpoint mechanism, the spindle assembly checkpoint which blocks mitotic exit and the next round of DNA replication upon spindle disruption, is weak or entirely absent in *Chlamydomonas* [35].

### Complementation of temperature-sensitive lethal mutations with fluorescent transgenes

We generated universal plasmids containing fluorescent tags, into which we inserted wild-type sequences to rescue the corresponding ts mutants. Using the rescued strains, we analyzed protein expression and localization (Figure 7, 9 and 11). We used *mCherry* tagged strains for indirect immunofluorescence because anti-mCherry antibody generally works better than anti-GFP antibody. For time-lapse microscopy, mCherry signal is hard to detect due to chloroplast autofluorescence in *Chlamydomonas*, but cell-cycle-regulated accumulation and localization of MCM4-Venus was clearly detectable at high time resolution (Figure 8).

### A loss of nuclear retention in a “closed mitosis”

Whether the nuclear envelope breaks down in mitosis or remains intact (‘open’ vs. ‘closed’ mitosis) is a significant phylogenetic marker, with multiple independent transitions in different lineages [53]. *Chlamydomonas* is traditionally classified as having a closed mitosis; however, permeability of the nuclear envelope in a closed mitosis may vary among organisms [54]. In *Chlamydomonas*, MCM4 and MCM6 were localized to the nucleus throughout the division cycle except during mitosis, when they transiently diffused to the cytoplasm; CDC6 may show a similar localization pattern (Figure 7A, 8A, 9C). Transient MCM protein diffusion during mitosis is also observed in *Arabidopsis* [31], which has an open mitosis.

*Chlamydomonas* spindle poles are outside the nuclear envelope and the spindle enters the nucleus through large holes called polar fenestrae [55]. The fenestrae are 300-500um in diameter, 5 times larger than nuclear pores in interphase. If the fenestrae are truly open channels, this should allow rapid diffusion of all nuclear components that are not tightly bound to large structures (such as chromosomes, mitotic spindles or a nuclear matrix). This transient protein diffusion might therefore be functionally equivalent to nuclear membrane breakdown in organisms showing ‘open mitosis’. Nuclear envelope breakdown is triggered in early mitosis and nuclei are reassembled at the end of mitosis. In an early proposal to explain ‘licensing’ of replication origins in *Xenopus*, it was proposed that the nuclear envelope excludes access of key replication factors to fired origins; nuclear envelope breakdown allows access and subsequent origin licensing [56]. We speculate that the fenestrae in *Chamydomonas* may serve as a related cell cycle reset mechanism. Our results suggest that the classic distinction between ‘open’ and ‘closed’ mitosis may not imply restricted traffic of nuclear components [54]. A general loss of nuclear integrity, rather than a specific effect on replication proteins, is suggested by transient diffusion of ble-GFP from the nucleus during mitosis (see above).

In contrast, CDC45-mCherry was stably nuclear throughout the cell cycle. In metaphase, when other proteins diffused through the cytoplasm, CDC45 appeared colocalized to spindles (Figure 11B); this anchoring may prevent CDC45 cytoplasmic diffusion in metaphase. We do not yet know the functional significance of CDC45 spindle localization.

### Pre-RC formation in *Chlamydomonas*

All of the DNA replication proteins we analyzed were essentially absent during the long G1 before multiple fission (Figure 6). This is correlated with transcriptional profiles of corresponding genes [5, 6]. In yeast, CDK activity is very low specifically in late mitosis and early G1, and low CDK activity is permissive for pre-RC formation and origin loading with the MCM helicase; loaded origins persist through G1 until firing occurs under positive CDK control. The essential absence of pre-RC proteins in *Chlamydomonas* during G1 suggests that origin loading may be regulated differently than in yeast, since during most of G1 there appears to be negligible levels of pre-RC proteins; however, we lack a direct origin loading assay to pursue this at present. In yeast, such studies were greatly facilitated by specific origin sequences (ARS consensus). We do not know if *Chlamydomonas* has specific origin sequences, or, by contrast, if origin selection is largely sequence-nonspecific as in animal cells.

In yeast, CDK phosphorylates the MCM complex, which triggers its nuclear export during S-phase after individual origins have fired. Our results show that MCM6 stays in the nucleus in *cdka1* and *cdkb1* mutants (Figure 7), which could be consistent with this mechanism. However, in wild-type cells MCM4 stays in the nucleus throughout the cell cycle except for a window of ~3 min, indicating that CDK probably does not exclude MCM4 from the nucleus during S-phase in *Chlamydomonas*, in contrast to results in yeast.

CDC6 also stayed in the nucleus in the *cdkb1* mutant (Figure 9). However, CDC6 was undetectable in the *cdka1* mutant (Figure 9 and 10). This was a surprising result because Cdc6 degradation is mediated through CDK-dependent phosphorylation in yeast. *CDKA1* expression peaked during S-phase 1 hour before *CDKB1* comes up [5, 6]. Efficient and timely expression of DNA replication genes depends on CDKA1 [5]. The Cdc6 suppression in *cdka1* mutant might be a reflection of this transcriptional inhibition. Cdc6 regulation is diverse across the species. In budding yeast, CDK mediates SCF-dependent Cdc6 degradation to inhibit pre-RC formation in S-phase. In humans, APC-dependent Cdc6 proteolysis inhibits Cdc6 accumulation, and CDK phosphorylates and protects Cdc6 to trigger origin licensing [23]. It is unclear whether any of these mechanisms will apply in *Chlamydomonas*. Proteasome-dependent degradation of CDC6 may provide a first clue (Figure 10B).

*Chlamydomonas* ts-lethal mutations in many proteins controlling DNA replication allow a detailed functional examination of DNA replication control in the plant kingdom. Absence of an accessible culture system or conditional alleles, and a high level of gene duplication makes such experimentation (and even identification of the cognate proteins; see above) much more difficult in land plants. As expected from phylogenetic relationships, conserved *Chlamydomonas* proteins are more closely related to land plant proteins than to the orthologous yeast or animal proteins (Figure 2). This closer conservation likely extends to functional considerations as well; thus *Chlamydomonas* may be a valuable model for replication control in the critically important plant kingdom.

The budding yeast DNA replication control system has been characterized in depth and detail, providing specific molecular hypotheses to evaluate in *Chlamydomonas*. The results reported here already demonstrate conserved features but also some likely and significant differences. It is important to note that conservation of sequence alone does not guarantee conservation of function or regulation. Conditional mutants are key resources to rapidly deliver functional analysis.

Sequence comparisons indicate that in some cases, *Chlamydomonas* and human replication proteins are much more homologous than the corresponding yeast and human proteins are (for example, DIV20/RecQL4/SLD2 discussed above). This is a genomic feature applying to many conserved proteins and presumably reflects rapid sequence evolution in the yeast lineages [57]. Whereas *Chlamydomonas* molecular evolution is quite slow, leading to retention of sequences and features found in animals but not in yeast [58]. Thus, although clearly in its infancy compared to the highly developed yeast systems, study of *Chlamydomonas* replication may provide useful models for features found in animals but lost in yeast.

## MATERIALS AND METHODS

### Cell culture conditions

All strains are listed in Table S1. Cells were maintained on Tris-acetate-phosphate (TAP) medium (Harris, 2008) at 21°C under continuous illumination at ~100 PAR (determined with Apogee Quantum Light Meter). In order to synchronize cells in G1 by nitrogen depletion, cells were spread on 0.1xN TAP plates (containing 1/10 of standard TAP levels of ammonium chloride) and incubated for 2 days at 21°C under the light. The G1 arrest was released by plating cells on TAP plates at 33°C under continuous illumination at ~150 PAR.

### Strain cross and mutant analysis

Strain cross was performed as described previously [59]. Strain list is in Table S1. Tetrad haploid progenies were dissected with a Zeiss Axioskop 40 Tetrad microscope. Mutations were verified by allele-specific competitive PCR (Onishi M). Briefly, three primers were designed for each mutant. Two forward primers were designed to distinguish wild type and mutant by a length of PCR product. Forward primer 1 contains wild type sequence with 20bp long. Forward primer 2 contains the mutant sequence with 40bp long. A reverse primer was designed to recognize sequence a few hundred base pairs downstream of the mutant site. PCR reaction was performed using these 3 primers, and the PCR product was analyzed using 3% SB agarose gel. The PCR band with the mutation is 20bp longer than that with wild type. Primer sequences are listed in Table S2.

### Plasmid construction and transformation

The mCherry plasmid *FC1011* plasmid was constructed by NEBuilder HiFi DNA assembly Master Mix (New England Biolabs, MA) using pKA1 (*CDKB* plasmid) as a temperate [41]. The mCherry fragment was replaced by GFP and Venus to create GFP- and Venus-universal plasmids, named *FC1012* and *FC1001* using multiple cloning sites. The desired gene (*MCM6, CDC45*, or *CDC6*) was amplified from genomic DNA and inserted into the mCherry plasmid at the multiple cloning site. *ORC1-mCherry* and *MCM4-VENUS* plasmids were synthesized by SynBio (NJ). For these plasmids, all of the introns except the first two and the last intron were removed. The plasmid linearized by XbaI digestion was used for transformation.

Temperature sensitive mutant cells were transformed with the linearized plasmid by electroporation (500 V, 50 μF, 800□Ω) using GenePulser Xcell (Bio-Rad, CA). The cells were recovered in TAP containing 40□mM sucrose, incubated under light overnight, and then plated on a TAP plate containing 10 ug/ml paromomycin. Complementation of temperature sensitivity among paromomycin-resistant transformants was determined by serial dilution of cells on TAP plate at 33°C for 2-5 days. Plasmid integration and structure was confirmed by PCR.

### Flow Cytometry

Flow Cytometry analysis (FACS) was performed as described by BD Accuri C6 instrument (BD Biosciences)[35]. The data was analyzed by FlowJo software (FlowJo, LLC, OR).

### Western Blotting

Cells were lysed in RIPA buffer containing protease inhibitors with glass beads using FastPrep (MP Biomedicals, CA) for 20s, twice, at speed 6. Cleared cell lysate was denatured with SDS buffer and separated by SDS–PAGE with Novex 4–12% Tris-glycine polyacrylamide gel (Invitrogen, Life Technologies, CA). Western blot analysis was performed using anti-mCherry antibody (16D7) at 1:2000 dilution (Thermo Fisher, MA), anti–GFP antibody (GF28R) at 1:2000 dilution (Thermo Fisher, MA) and anti-AtpB antibody at 1:5000 dilution (Agrisera, Sweden) as a loading control. HRP-conjugated secondary antibody was used to detect signal with SuperSignal West Femto (Thermo Fisher, MA). Image was acquired with Image Quant LAS 4000 (GE Healthcare, IL). The band intensity was quantified with ImageJ (NIH).

### Indirect Immunofluorescence

Indirect immunofluorescence was performed as described [35]. The primary antibodies are anti-mCherry monoclonal antibody (16D7) at 1:200 dilution (Thermo Fisher, MA) and anti-α-tubulin antibodies (clone B-5-1-2, Sigma-Aldrich) at 1:5000. The secondary antibody was anti-rat-Alexa 568 (Thermo Fisher, MA) at 1:1000 and DyLight™ 405 goat anti-mouse antibody (BioLegend, CA) at 1:2000, respectively. Cells were stained with 1 μM or 0.5 μM Sytox Green (Invitrogen, CA). Images were collected using an inverted Zeiss LSM 510 Laser Scanning confocal microscope. Images in Figure S1 were collected using DeltaVision microscope. Images were exported in TIFF format, merged and analyzed using ImageJ (NIH).

### Time-Lapse Microscopy

Cells arrested in G1 by nitrogen deprivation were transferred to agarose pads and imaged with a 63X objective on a Leica DMI6000B microscope with the objective and stage heated to 33°C. Pad architecture and other procedures were based on the budding yeast setup of Di Talia et al, but with modifications for *Chlamydomonas* to be described elsewhere (KP and FC, unpublished data)[46]. Images were acquired using custom software, as previously described for yeast timelapse [47], modified to adjust autofocus for *Chlamydomonas* (KP and FC, unpublished). Deconvolution was employed to subtract the chloroplast-specific signal in the Venus channel (KP and FC, unpublished). All images in a given movie were analyzed identically.

## SUPPLEMENTAL DATA

The following materials are available in the online version of this article.

**Supplemental Table 1**. *C. reinhardtii* strains used in this paper.

**Supplemental Table 2**. Oligonucleotides sequence used to verify the mutation. Lowercase letters indicate additional external sequence.

## ACKNOWLEDGMENTS

We thank Daria Postavnaia for help with making plasmids in Figure 4A. We also thank Alison North at The Rockefeller University Bio-Imaging Resource Center for technical assistance with confocal microscopy, and The Rockefeller University Flow Cytometry Resource Center for FACS analysis. This work was supported by NIH 5SC1GM121242 to AI and 2R01GM078153 to FC.

